# A small family business: synergistic and additive effects of the queen and the brood on worker reproduction in a primitively eusocial bee

**DOI:** 10.1101/756692

**Authors:** Margarita Orlova, Jesse Starkey, Etya Amsalem

## Abstract

The mechanisms that maintain reproductive division of labor in social insects are still incompletely understood. Most studies focus on the relationship between adults, overlooking another important stakeholder in the game – the juvenile offspring. Recent studies from various social species show that not only the queen, but also the brood regulates reproductive division of labor between females, but how the two coordinate to maintain reproductive monopoly remained unexplored.

Our study aims at disentangling the roles of the brood and the queen in regulating worker reproduction in primitively eusocial bees. We examined the effects induced by the brood and queen, separately and together, on the behavioral, physiological and brain gene expression of *Bombus impatiens* workers. We found that young larvae induce a releaser effect in workers, decreasing egg laying and aggressive behaviors, while the queen induces both releaser and primer effects, modifying worker aggressive and egg laying behavior and reproductive physiology. The expression of reproduction- and aggression-related genes was altered in the presence of both queen and brood, but the effect was stronger or the same in the presence of the queen.

We identified two types of interactions between the queen and the brood in regulating worker reproduction: (1) synergistic interactions regulating worker physiology, where the combined effect of the queen and the brood was greater than each of them separately; (2) additive interactions regulating worker behavior, where the combined effects of the queen and the brood are the gross sum of their separated effects. In these interactions the brood acted in a manner similar to the queen but to a much smaller extent and improved the quality of the effect induced by the queen. Our results suggest that the queen and the brood of primitively eusocial bees coordinate synergistically, additively, and sometimes even redundantly to regulate worker behavior and reproduction, and the interaction between them exists in multiple regulatory levels.

## Introduction

Reproductive division of labor is the defining feature of insect sociality. It exists in a variety of forms across multiple species, ranging from a modest reproductive skew by a few dominant, morphologically-identical females, to a complete monopolization of reproduction by a single queen (Wilson, 1971). However, the understanding of both the proximate and the ultimate causes of reproductive division of labor is incomplete. Pheromonal signaling and behavioral interactions are considered the most common mechanisms used by the colony members to enforce reproductive monopoly (Kocher and Grozinger, 2011). However, the different parties regulating worker reproduction and their relative roles remain largely unexplored.

Most proximate studies examining the regulation of reproduction have focused on interactions between adult members of insect societies, overlooking the potential role of juveniles. In a variety of species, queen’s behavior and pheromonal signaling, as well as interactions between nestmate workers, were found to affect worker reproduction (Ronai et al., 2016; Wenseleers et al., 2004). Queen pheromones have been identified in a small number of species (Hefetz, 2019; Le Conte and Hefetz, 2008) and possible mechanisms of their action are still debated (Keller and Nonacs, 1993; Smith and Liebig, 2017; Villalta et al., 2018), but queens and workers are not the only parties to the conflict over reproduction in social societies. Juveniles also hold stakes in the matter and are involved in reproductive conflict with other juveniles and adults (Ebie et al., 2015; Schultner et al., 2017; Starkey et al., 2019a; Starkey et al., 2019b; Ulrich et al., 2016a).

The main point of contention between juvenile and their caregivers lies in the fact that offspring are selected to demand more parental investment than parents are selected to provide (Trivers, 1972), resulting in a conflict over resource allocation between current brood and future generations (inter-brood conflict). This tradeoff is not unique to social animals and is well documented in both non-social vertebrates (Calisi et al., 2016; Weir and Rowlands, 1973), and invertebrates (Schultner et al., 2017). However, it is often overlooked in social insect studies, that are centered around the role of royals in shaping the social structure of the colony. One exception is the honey bee *Apis mellifera* where the role of brood was extensively examined, showing that pheromones produced by the brood regulate worker reproduction and maturation (Maisonnasse et al., 2010; Maisonnasse et al., 2009; Mohammedi et al., 1998). Several recent studies further highlight the role of brood in regulating worker reproduction and behavior in ants (Ebie et al., 2015; Ulrich et al., 2016b) and bumble bees (Starkey et al., 2019a). However, even in these species, the interplay between the roles of juveniles and adults remained understudied, partly because in eusocial insect societies, queen and brood exert their influence on workers simultaneously, and the effects of the queen and the juveniles are difficult to disentangle.

Derived eusocial species are less informative about the mechanisms regulating reproduction since, in many cases, they reached ‘a point of no return’ where worker sterility can no longer reversed. The bumble bee *Bombus impatiens* is an excellent model system to study the effects of brood and queen on reproductive division of labor, since they are a primitively eusocial species with relatively small colonies, limited morphological differences between castes (Amsalem et al., 2015a; Michener, 1974) and high rates of worker reproduction (Alaux et al., 2004; Cnaani et al., 2002). Previous studies in bumble bees show that the queen inhibits worker reproduction during the first part of the social life cycle, but loses the ability to exert reproductive dominance later on during the ‘competition phase’, where the workers and the queen compete over male production (Cnaani et al., 2002; Duchateau and Velthuis, 1988; Padilla et al., 2016). Various chemical signals are produced by the queen or found in the wax, and although found to correlate with the queen’s fecundity in several cases (Amsalem et al., 2014a; Amsalem et al., 2015b; Rottler et al., 2013; Sramkova et al., 2008), are insufficient to inhibit worker reproduction independently from a freely behaving queen (Amsalem et al., 2015b; Amsalem et al., 2017; Padilla et al., 2016; Rottler-Hoermann et al., 2016; Van Oystaeyen et al., 2014).

Recent findings suggest that also the brood plays a role in the inhibition of worker reproduction in *B. impatiens* (Starkey et al., 2019a; Starkey et al., 2019b). Young, but not old larvae, reduced egg laying but not ovary activation in workers in a quantity-dependent manner, with nearly complete suppression of egg laying in groups containing two workers and 10 young larvae (Starkey et al., 2019a). This effect is unlikely to be solely mediated via pheromones and, similar to the queen’s impact on workers, requires a physical contact between the workers and the brood (Starkey et al., 2019b). The larval effects were independent of relatedness between the workers and the brood, brood sex or worker age, with both newly emerged workers or random-age workers showing the same pattern of response in the presence of brood (Starkey et al., 2019a). Larvae and pupae effects were examined using encased brood, however the wax itself or its extracts, although found to reduce ovary activation and aggression in small queen-right *B. terrestris* workers (Rottler-Hoermann et al., 2016), had no effect on worker reproduction in *B. impatiens* (Starkey et al., 2019a; Starkey et al., 2019b). Overall, both the brood and the queen affect *B. impatiens* worker reproduction, however, the respective roles of the brood and the queen and how they interact to regulate worker reproduction remained unresolved.

Our study endeavors to examine this question by studying the effects of the queen and the brood on worker reproduction at multiple regulatory levels, including worker reproductive physiology, egg laying and brood care behavior, aggressive behavior, and brain gene expression. In the first experiment we grouped pairs of workers with a queen, brood, both or none, and examined their effect on worker oocyte size and egg laying behavior. In the second experiment, we allowed pairs of workers to directly or indirectly interact with a queen, brood or both, and measured their aggressiveness and brood care behaviors. In the last experiment, we grouped pairs of workers with different types of brood (pupae, larvae, wax or none) or with a queen, brood, both or none, and measured the expression levels of four candidate genes in worker brains. All genes were previously found to regulate reproduction and/or aggression in bumble bee workers. We analyze the interactions between the queen and the brood in regulating worker reproduction and discuss possible mechanisms of reproductive regulation at different regulatory levels and by different players.

## Methods

### General bumble bee rearing

Colonies of *B. impatiens* were obtained from Koppert Biological Systems (Howell Michigan, USA), maintained in the laboratory under constant darkness, a temperature of 28–30°C, 60% relative humidity, and supplied *ad libitum* with a sugar solution and fresh pollen (Light spring bee pollen, 911Honey). These colonies were used as a source of callows (newly emerged workers <24 h) and brood. In all experiments, workers (n=346) were separated from their parental colonies and placed in pairs (n=173) in small plastic cages (11 cm diameter x 7 cm height) with different combination of brood and queen as compared to controls. Active egg laying queens were taken from full-size Koppert colonies.

#### Experiment 1 The effects of brood and queen presence on worker reproduction

Newly emerged workers were sampled from two parental colonies and placed in pairs for 10 days in order to allow them to fully activate their ovaries and lay eggs. Cages were randomly assigned to one of five treatments: (1) 8 pairs of workers without a queen or brood (w/o QB). Eggs that were laid by these workers (typically within 8-9 days) were counted and removed daily to maintain constant absence of brood; (2) 8 pairs of workers with 10-20 young larvae (B). Clutches of 10-20larvae encased is a thin wax envelope separated from other wax structures were placed in the cages at the onset of the experiment and allowed to develop normally. The feeding period of *B. impatiens* larvae lasts 9-11 days (Cnaani et al., 2002), thus all larvae turned into pupae by the end of the experiment. Eggs that were laid in these cages remained untouched and were counted by the end of the experiment; (3) 8 pairs of workers with 10-20 young larvae (as described earlier) were replaced five days after the onset of the experiment with similar amount of new young larvae (YB). In a previous study we found that only young larvae reduce worker egg laying while pupae induce the opposite effect (Starkey et al., 2019a). Therefore, this procedure ensured constant presence of young larvae throughout the experiment. Eggs that were laid in these cages remained untouched and were counted by the end of the experiment; (4) 9 pairs of workers with a queen but without brood (Q). Eggs that were laid in these cages (typically by the queen within 1-2 days) were counted and removed daily to maintain constant absence of brood; (5) 9 pairs of workers with a queen and 10-20 young larvae (QB). Eggs that were laid in these cages (typically by the queen within 1-2 days) remained untouched and were counted by the end of the experiment. Diagram of the experimental design is provided in Fig 1a. All cages were kept for 10 days, after which workers were frozen at −20° C until further analysis. We collected data about worker and queen egg laying and worker oocyte size.

**Fig. 1A.**
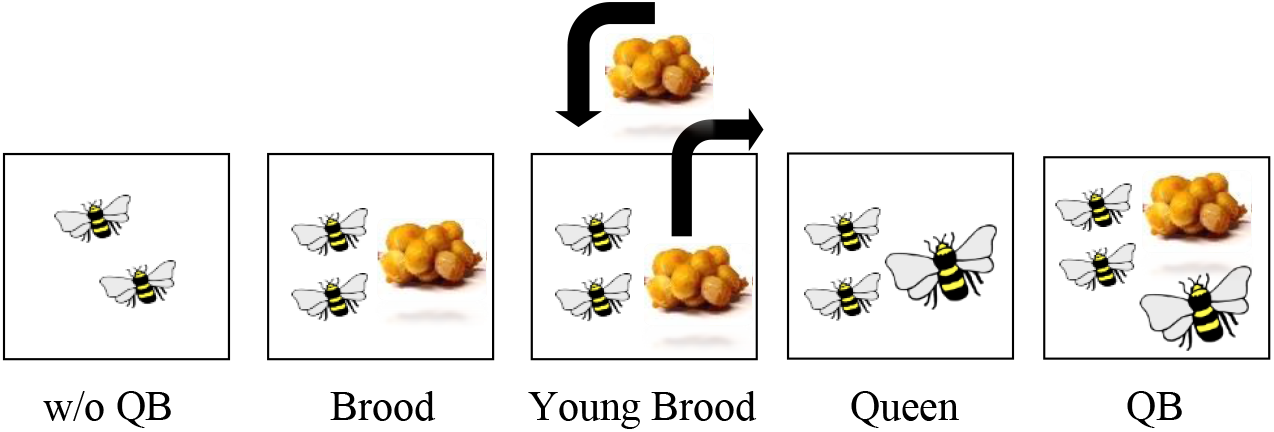
Diagrams of the experimental design of experiment 1

#### Experiment 2 The effects of brood and queen presence on worker aggressive and brood care behaviors

In this experiment we tested the effects of brood and queen on worker behavior and also the effects they may have on worker behavior when perceived indirectly through a mesh. Newly emerged workers were collected from four parental colonies and housed in a rectangular cage divided in two by a mesh screen for 3 days. Previous studies show that the majority of aggression is exhibited by workers within 3 days (Amsalem and Hefetz, 2010; Padilla et al., 2016). In each compartment we placed a pair of workers that was randomly assigned to one of the following treatments: (1) direct contact with 10-20 young larvae (12 pairs); (2) indirect contact (through a mesh) with 10-20 young larvae (12 pairs); (3) direct contact with an active queen but without brood (11 pairs); (4) indirect contact with an active queen without brood (11 pairs); (5) direct contact with an active queen and 10-20 young larvae (12 pairs), and (6) indirect contact with an active queen and 10-20 young larvae (12 pairs). Diagram of the experimental design is provided in Fig 1b. All the eggs found in the cages were laid by the queen. These eggs remained untouched or were counted and removed daily in compartments that were designed to remain brood-less. Observations were carried out for 20 minutes per pair per day during days 1-3. Observations were conducted daily between 12:00 to 16:00. During observations we recorded aggressive interactions between workers in each pair and interactions between adult workers and brood. Aggressive interactions included climbing (one bee mounting another bee), humming (rapid wing movements directed at another bee without a physical contact), darting (rapid movement towards another bee without a physical contact), pushing (physical contact from which the other bee retreats) and attack (overt fight with biting and stinging attempts), as described in (Amsalem and Grozinger, 2017; Amsalem and Hefetz, 2010). All these behaviors are performed in a higher rate by dominant bumble bee females, both workers and queens (Amsalem and Grozinger, 2017; Amsalem and Hefetz, 2010; Amsalem and Hefetz, 2011; Amsalem et al., 2014b; Amsalem et al., 2014c; Duchateau, 1989; Padilla et al., 2016). The sum of all aggressive behaviors that occurred during the observation period per cage was termed ‘aggression index’ and used in further analysis. Interactions with brood included feeding and incubation. The sum of all interactions with brood per cage was termed ‘brood-tending index’ and used in further analysis. Workers were sampled on the fourth day by flash freezing and kept in −20° C until further analysis.

**Fig. 1B.**
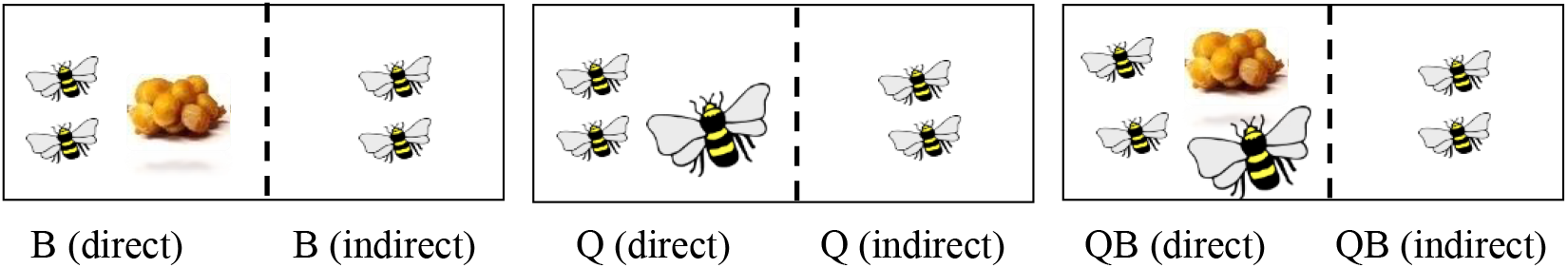
Diagrams of the experimental design of experiment 1

#### Experiment 3 The effect of queen and brood on worker brain gene expression pattern

Newly emerged workers were collected from four parental colonies and kept in pairs for 3 days to capture gene expression differences before workers activate their ovaries. Pairs of workers were randomly grouped with: (1) no brood or queen (9 pairs); (2) piece of wax (10 pairs); (3) 10-20 young larvae encased in a wax envelope (9 pairs), and (4) approximately 10 pupae encased in their individual cocoons envelopes (9 pairs). In a follow up experiment, we grouped newly emerged workers with (1) no brood or a queen (w/o QB, 6 pairs); (2) 10-20 larvae (B, 6 pairs); (3) an active queen with no brood (Q, 6 pairs). Eggs laid by the queen in these cages were counted and removed daily; (4) a queen and 10-20 larvae (QB, 6 pairs). In these cages, the queen’s eggs remained in the cage and were counted by the end of the experiment. Diagrams of the experimental design are provided in Figs 1c and 1d. Workers were sampled on the fourth day by flash freezing and kept in −80° C until further analysis. We extracted RNA from worker brains (pool of 2 brains from the same cage per sample) and collected data about worker oocyte size and brain gene expression.

**Fig. 1C.**
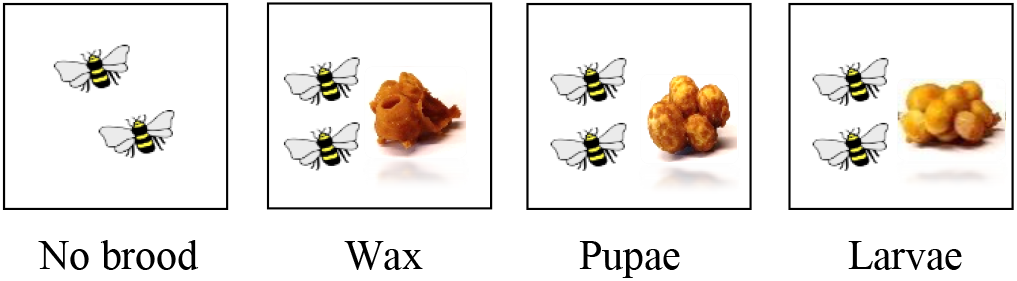
Diagrams of the experimental design of experiment 3a

**Fig. 1D.**
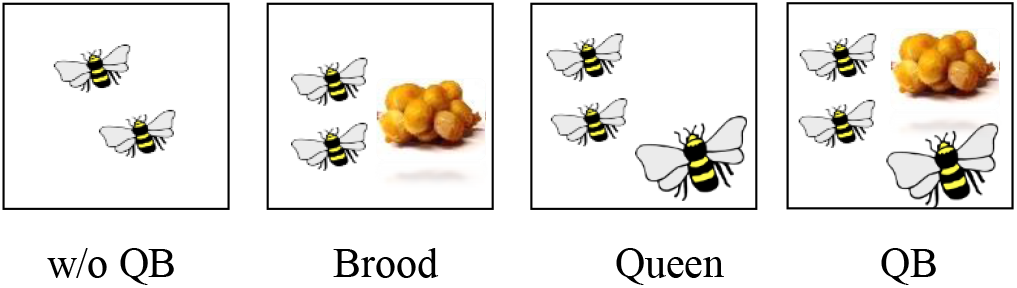
Diagrams of the experimental design of experiment 3b

### Brood and wax collection

Larvae, pupae and wax were gently removed from their parental colonies and were used only if they remained intact during collection. Young larvae were defined by their mass (<50 mg, roughly corresponding to instars 1 and 2) as in our previous studies (Starkey et al., 2019a; Starkey et al., 2019b). All the brood and the wax were collected from queen-right colonies with no signs of worker reproduction. All the brood was likely to be of female workers, though we have previously shown that worker egg laying is similarly decreased regardless of the brood sex (Starkey et al., 2019a). In all experiments, workers were introduced to unrelated brood. In a previous study we showed that worker reproduction is similarly affected by related or unrelated brood (Starkey et al., 2019a).

### Egg laying

Egg laying by queens and workers was observed daily. The cumulative number of eggs (or larvae, if eggs hatched) was counted by the end of each experiment. While egg oophagy generally exists in bumble bees, it is often performed in queen-right colonies and rarely occurs in small queen-less groups (Amsalem et al., 2015a). We did not see evidence for oophagy (such as open egg cells, etc.) that could affect the results. To account for variation in worker egg laying between colonies (Amsalem et al., 2015a; Amsalem et al., 2015b), we ensured that each experiment was replicated using several source colonies, equally representing both treatment and control groups. We statistically controlled for colony effect whenever such effect was found.

### Measurement of ovarian activation

After bees were collected, each bee was placed in a separate tube and received an individual number corresponding with their cage and treatment. Thus, dissections were performed blindly. Ovaries were dissected under a stereomicroscope and placed into drops of distilled water. The length of the terminal oocyte in the three largest ovarioles was measured with a micrometer eyepiece embedded into the lens. Workers possess four ovarioles per ovary and at least one oocyte per ovary was measured. Mean terminal oocyte length for each bee was used as an index of ovarian activation (Amsalem et al., 2009)

### Brain dissections and RNA extraction

Bumble bee workers were collected by flash freezing on dry ice. Heads were separated from the thorax and stored at −80°C until RNA extraction. Brains of each pair of bees were separated from the head on dry ice and pooled together. Total RNA was extracted using a RNeasy kit (Qiagen) according to the manufacturer’s instructions. RNA quality and quantity were analyzed using a NanoDrop One^C^ (Thermo Scientific).

### Primer design and choice of genes

Genes were identified using the NCBI/blast home page (http://blast.ncbi.nlm.nih.gov/Blast.cgi). Design of forward and reverse primers for each gene was performed using PrimerBLAST https://www.ncbi.nlm.nih.gov/tools/primer-blast/) or was taken from a previous study (Padilla et al., 2016). A list of all primers used in this study is provided in Table S1. Four genes have been selected based on previous studies showing that they are regulated in bumble bees in association with reproduction, aggressive behavior or both: (1) *Vitellogenin (vg)* is the major egg yolk protein female insects invest in the ovaries (Hagedorn and Kunkel, 1979). Vg was upregulated in aggressive and fertile workers (vs. subordinate and sterile) and queens (vs. workers) of *B. terrestris* fat-body and heads (Amsalem et al., 2014b), and downregulated in the presence of the queen in *B. impatiens* heads (vs. her absence) (Padilla et al., 2016). Brain *vg* levels were shown to differentially expressed in *B. terrestris* as function of sex, caste and reproduction (Jedlicka et al., 2016b); (2) *Kruppel homolog 1 (krh1)* is a transcription factor upstream to the juvenile hormone (JH) synthesis and was upregulated in dominant *B. terrestris* workers (vs. subordinates) and in the absence of the queen (vs. its presence) (Shpigler et al., 2010), but did not decrease in workers in the presence of the queen in *B. impatiens* (Padilla et al., 2016) and was upregulated in both diapausing queen and males of *B. terrestris* (Jedlicka et al., 2016a); (3) *Methyl farneosoate epoxidase (mfe)* encodes to the final enzyme in the synthesis of JH and was downregulated in non-reproductive *B. terrestris* queens (vs. reproductives) (Jedlicka et al., 2016a); (4) *DNA methyltransferase 3 (dnmt3)* encodes to the DNA methyltransferase enzyme that is essential for creating de novo DNA methylation marks on the genome and was upregulated in older *B. terrestris* workers (vs. younger) (Lockett et al., 2016). It was also associated with reproductive castes in the honeybee (Kucharski et al., 2008). However a recent study found no evidence for methylation directly affecting gene expression between reproductive and sterile workers in *B. terrestris* (Marshall et al., 2019).

### Gene expression analysis

Synthesis of cDNA (Applied Biosystems™) was performed according to the manufacturer’s instructions using 200 ng of RNA. Two μl of diluted cDNA were combined with 5μL SYBR-Green Master mix (Bioline, Luckenwalde, Germany), 0.2 μl of each forward and reverse primer (10 ɥM stock) and 4.6 μl DEPC-water. Two housekeeping genes were used to control for PCR efficiency: Arginine kinase and Phospholipase A2. These genes were found to be stable in *B. impatiens* brains and were used in several of our previous studies (Amsalem and Grozinger, 2017; Padilla et al., 2016). Expression levels were determined using qRT-PCR on a QuantStudio 5. Negative control samples (cDNA reaction without RT enzyme) and a water control were also present on each plate. PCR product quality and specificity were verified using melt curve analysis. Triplicate reactions were performed for each of the samples and averaged for use in statistical analysis. Expression levels of candidate genes were normalized to the geometric mean of two housekeeping genes using the 2^-ΔΔCt^ technique.

### Statistics

Statistical analyses were performed using SPSS v.21. Generalized Estimating Equations analysis (hence GEE) was employed for all comparisons. The models were built to control for interdependencies within data using parental colony and cage as subject variables. Worker ID and direct/indirect contact were used as a within-subject variables for oocyte size and behavior analyses respectively. Poisson loglinear distribution was used for egg laying analysis. Unstructured correlation matrix was used in models for egg-laying, oocyte size and behavior analysis. Exchangeable correlation matrix was used in models for gene expression analysis. Robust estimation was used to handle violations of model assumptions. All analyses used treatment as the main effect and were followed by post-hoc contrast estimation using Least Significant Difference (LSD) method. Non-parametric Spearman’s rho was used for correlation analyses. For genes that were significantly correlated with oocyte size, the latter parameter was used as a covariate in the GEE model analysis to control for its effect. Data are presented as boxplots featuring the minimum and maximum values, outliers and medians (egg laying, oocyte size, and aggressive behavior), or as means ± S.E.M. (gene expression). Statistical significance was accepted at α=0.05.

## Results

### Experiment 1 The effects of brood and queen presence on worker reproduction

Worker egg laying was the highest in the absence of queen or brood (w/o QB), significantly reduced in the presence of larvae that developed into pupae throughout the course of the experiment (B) and further reduced in the continuous presence of young larvae (YB). In both queen groups (Q, QB), no egg laying by workers was observed, regardless of the presence of brood (GEE, Wald χ^2^_2_= 33.63, p<0.001 for treatment, significant post-hoc contrasts indicated by different letters in Fig. 2a).

**Fig. 2:**
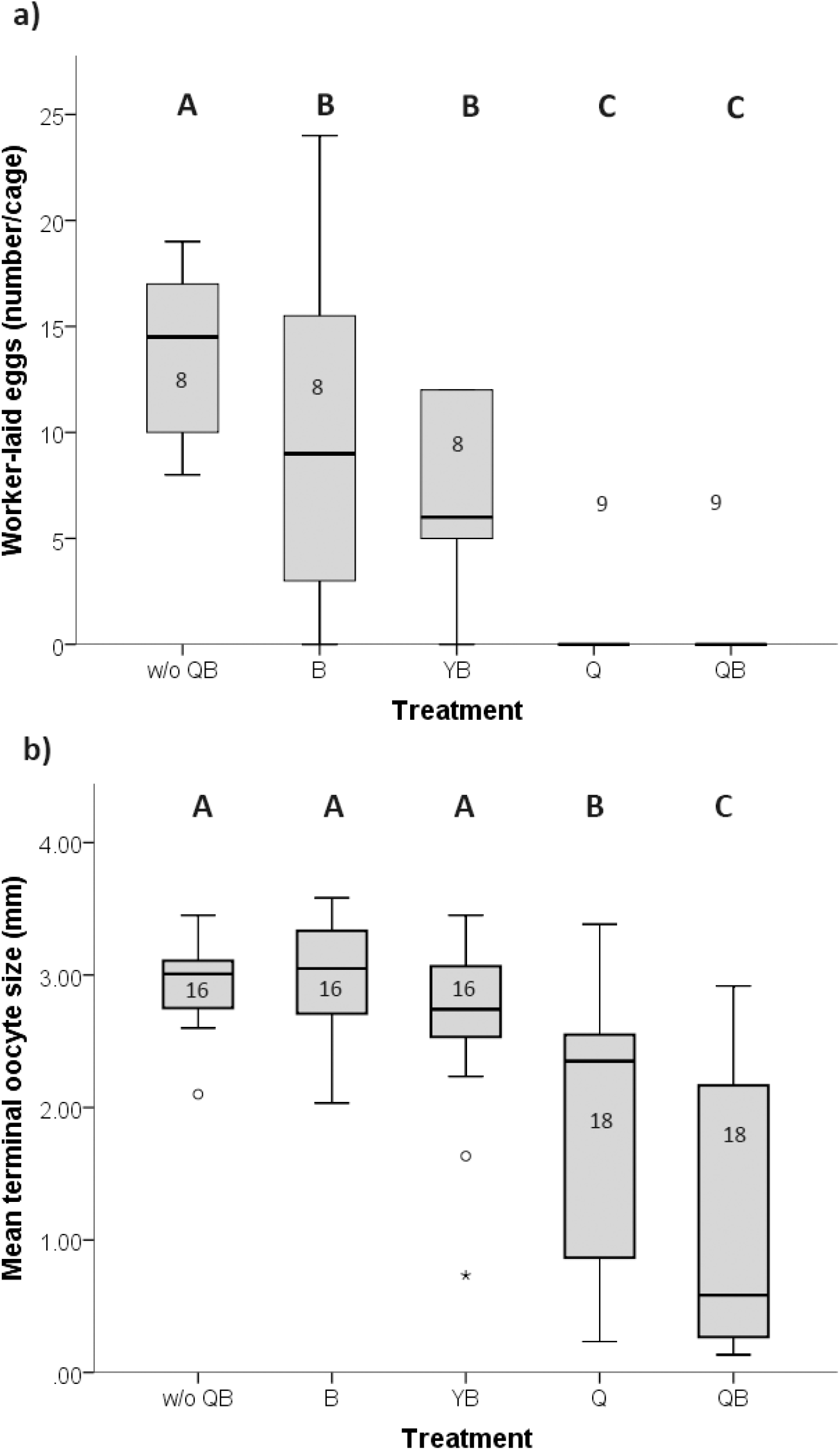
Number of worker-laid eggs (a) and oocyte size (b) across different treatments. Box plots display medians, quartiles and minimum and maximum values. Dots above/below each box indicate outliers. Statistical differences are reflected by different letters above boxes. Sample size is indicated within boxes.

We further examined the effect of the treatment on worker oocyte size. The three queen-less groups did not differ in oocyte size with all workers exhibiting fully activated ovaries. However, worker oocyte size was significantly reduced in the presence of the queen and even more so in the presence of the queen with brood (GEE, Wald χ^2^_4_= 58.45, p<0.001 for treatment, significant post-hoc contrasts indicated by different letters in Fig. 2b).

### Experiment 2 The effects of brood and queen presence on worker aggressive and brood care behaviors

Aggressive interactions between workers (see Methods) were counted for 20 minutes per day for three days and summed together for each cage. Levels of aggression between workers differed significantly across treatments and exposure types (direct/indirect) but the interaction between the two factors was not significant (GEE, Wald χ^2^_2_ = 10.97, p=0.004 for treatment, Wald χ^2^_1_ = 18.95, p<0.001 for exposure type, Wald χ^2^_2_ = 3.81, p=0.149 for interaction; Fig. 3).

**Fig. 3:**
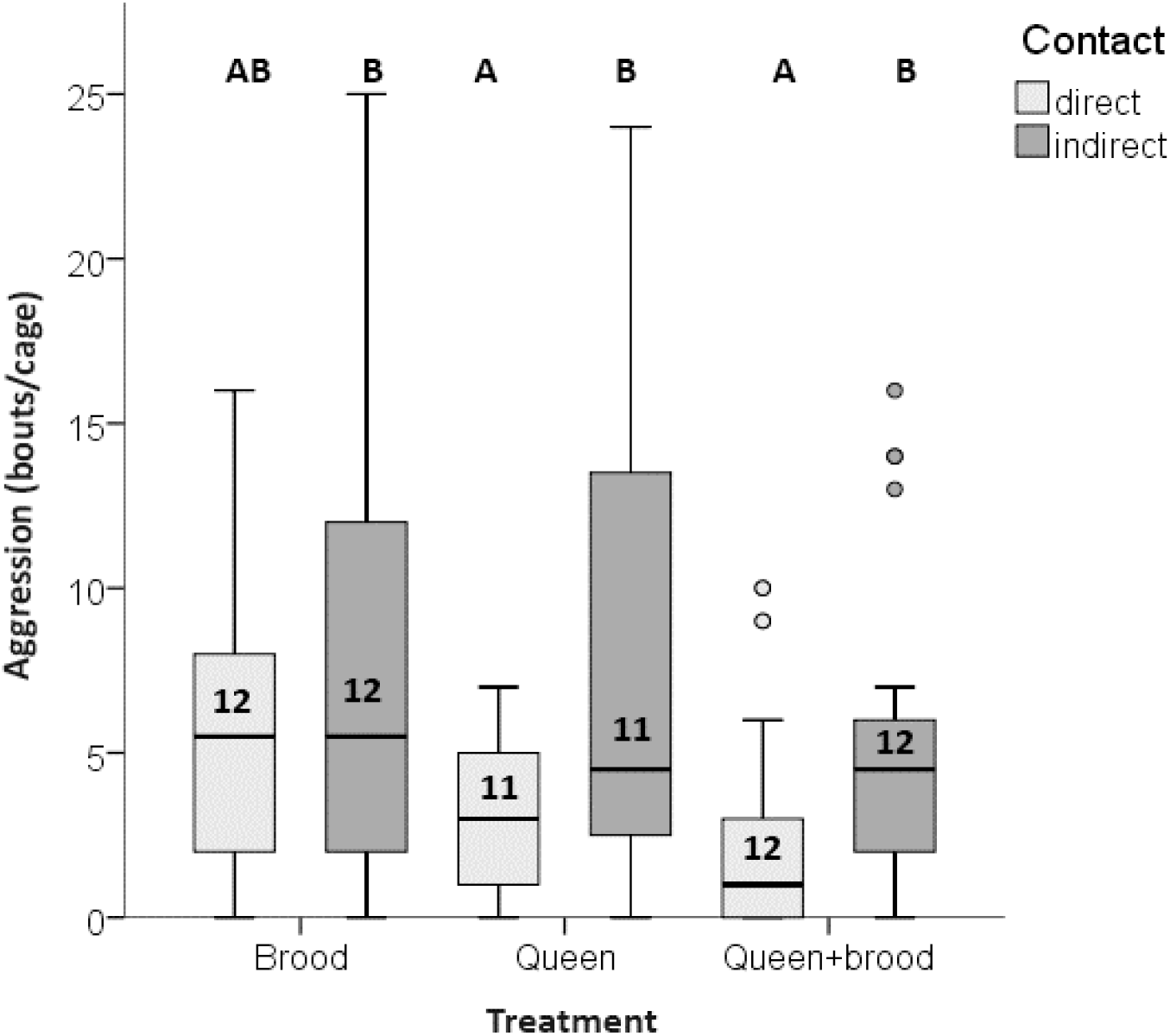
Number of aggressive interactions across different treatments with direct contact (light boxes) and indirect contact (dark boxes) with brood, queen or both. Box plots display medians, quartiles and minimum and maximum values. Dots above/below each box indicate outliers. Statistical differences are reflected by different letters above boxes. Sample size is indicated within boxes.

Worker aggression levels were significantly reduced only in the direct presence of the queen and even more so in the direct presence of the queen and the brood (post-hoc LSD, p=0.002 for queen and brood vs. brood alone, p=0.556 for brood alone vs. queen alone and p=0.013 for queen + brood vs. queen alone). The aggression levels of workers in the presence of the brood were intermediate compared to the queen-right groups and the controls (post-hoc LSD, p=0.087 for direct vs. indirect exposure to brood, p=0.556 for direct exposure to brood vs. queen).

Brood tending behaviors (the number of feeding and incubating events workers performed) were observed and counted in cages with direct exposure to brood and queen as compared to cages with only brood (i.e., the only cages where brood was present). The total number of brood-tending behaviors was greater in cages with queen and brood compared to cages with only brood (24±2.53 and 35.25±6.6 respectively, GEE, Wald χ^2^_1_ = 11.69, p=0.001). However, brood tending behaviors per capita (i.e., divided by the number of bees tending the brood in each cage, since together with the queen, queen-right cages included three females compared to only two in the queen-less cages) did not significantly differ between the queen-right and the queen-less groups (4.2±1.26 and 7.62±2.2 respectively, GEE, Wald χ^2^_1_ = 0.107, p=0.743).

### Experiment 3 The effect of brood type, and queen and brood on worker brain gene expression pattern

In the first experiment we kept pairs of newly emerged workers for 3 days with no brood, piece of wax, 10 pupae or 10-20 young larvae. At the time of sampling all these workers had inactive ovaries (0.22±0.01 mm, n=74). However, oocyte size significantly correlated with *krh1* expression levels (Spearman’s ρ=0.38, p=0.021) and therefore was included in the analysis as covariate.

*Vg* and *mfe* expression levels differed significantly across treatments (GEE, Wald χ^2^_3_ = 20.04, p<0.001 and GEE, Wald χ^2^_3_ = 12.02, p=0.007 respectively). *Vg* levels were significantly lower in pairs exposed to larvae compared with pairs kept without brood or wax (post-hoc LSD contrast, p=0.003 and p<0.001 respectively). *Mfe* levels were significantly lower in pairs exposed to larvae and pupae compared to pairs housed without brood (post-hoc LSD contrast, p=0.04 for both comparisons; Fig. 4). Expression levels of *krh1* did not differ significantly across treatments but covaried significantly with oocyte size (GEE, Wald χ^2^_3_ = 0.11, p=0.99 and GEE, Wald χ^2^_1_ = 4.182, p=0.041 respectively). *Dnmt3* expression also did not differ significantly across treatments (GEE, Wald χ^2^_3_ = 5.06, p=0.168; Fig. 4).

**Fig. 4:**
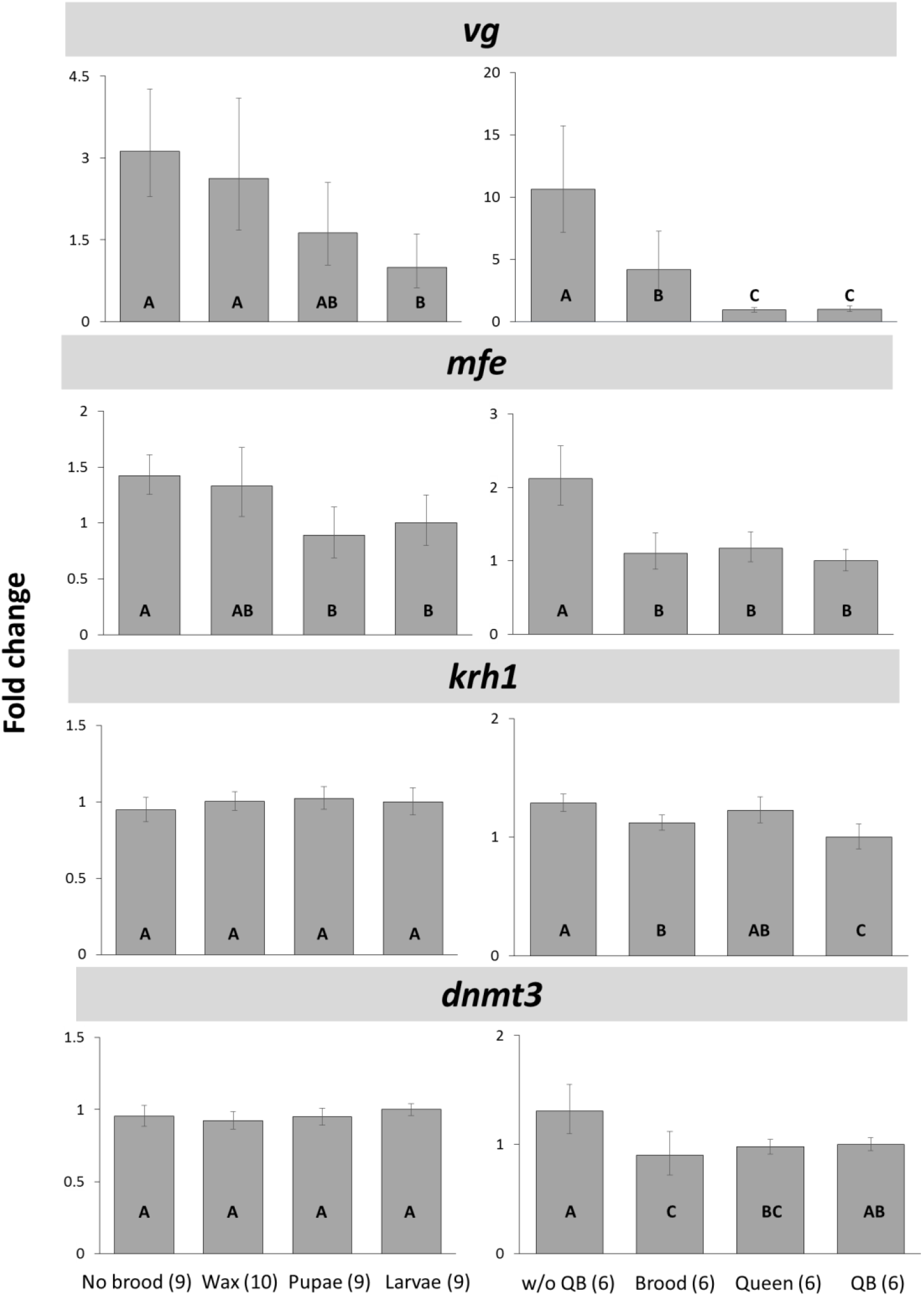
Gene expression levels across treatments with different brood types (a) and combinations of queen and brood exposure (b). Expression levels are displayed as fold changes relative to larvae treatment (a) and QB treatment (b). Bars represent mean fold change with error bars calculated from minimum and maximum Ct difference. Statistical differences are reflected by different letters within bars. Sample sizes for each treatment are indicated in parentheses.References

In the second experiment, we examined the effect of brood and queen presence on workers brain gene expression. Here too, workers were 3 days old at the time of sampling and all workers had inactive ovaries (0.2±0.01 mm on average, n=48 workers). However, oocyte size correlated significantly with *vg* expression (Spearman’s ρ=0.53, p=0.007). When *vg* expression was compared across treatments with oocyte size as a covariate, the difference between treatments was significant but the covariance with oocyte size was not (GEE, Wald χ^2^_3_ = 16.83, p<0.001 and GEE, Wald χ^2^_1_ = 3.01, p=0.08 respectively). Post-hoc comparisons revealed that pairs kept with brood, without brood and queen and in queen-right treatments (with or without brood) all differed significantly (post-hoc LSD, p<0.004 for all comparisons), but the queen-right treatments were similar (post-hoc LSD, p=0.67). *Mfe* expression levels also differed significantly across treatments (GEE, Wald χ^2^_2_ = 39.78, p<0.001) with pairs without queen and brood displaying higher expression levels compared to all other treatments, but the later three were similar to one another (post-hoc LSD, p<0.001 for control vs. all other treatments, p>0.3 for all other comparisons; Fig. 4). *Krh1* and *dnmt3* expression also differed across treatments (GEE, Wald χ^2^_2_ = 15.42, p<0.001 and GEE, Wald χ^2^_2_ = 12.234, p=0.002 respectively; Fig. 4). *Krh1* levels were highest in control groups and lowest in workers grouped with queen and brood, while with brood alone or queen alone the levels were intermediate. *Dnmt3* levels were highest in control groups and lowest in groups with brood alone, while queen-right treatments showed intermediate expression.

## Discussion

Our results offer meaningful insights into the effects of the queen and the brood on *B. impatiens* worker reproduction. We show that while the effect induced by the queen was always stronger than the brood, brood on its own or with the queen exerts a meaningful effect on worker reproduction and that this effect is manifested at multiple levels – from altering expression of genes in worker brain to decreasing egg-laying and aggressive behaviors. We have identified three different interactions between the brood and the queen roles in our data (Table 1): (1) ***synergistic effects***: neither the queen nor the brood alone were able to induce the full effect in workers but the combined effect of the brood and the queen was stronger than each of the effects alone; (2) ***additive effects***: the combined effects of the queen and the brood are the gross sum of their separated effect. In these interactions brood acted in a manner similar to the queen but to a much smaller extent and improved the quality of the effect induced by the queen; and (3) ***redundant effects***: the brood effect was equal to the effect induced by the queen, and either the brood or the queen were able to induce the same effect in workers.

**Table 1.** The regulatory interactions of the queen and the brood on worker reproduction in *B. impatiens*

| TYPE OF INTERACTION | PATTERN | EXAMPLES | REDUNDANCY | SHARED THEME |
| --- | --- | --- | --- | --- |
| SYNERGISTIC | QB>Q, Q>B | Oocyte size | No redundancy | Regulation of physiology |
|  | QB>Q, Q=B | *krh1* levels |  |  |
| ADDITIVE | QB=Q, Q>B | *vg* levels<br>Egg laying<br>Aggression | B is redundant to Q, but<br>Q is not redundant to B | Regulation of behavior |
|  | QB=Q=B | *mfe* levels<br>*dnmt3* levels | Q and B redundant | Unknown |

In a few cases we found that the queen and the brood acted on worker reproduction in synergy. The combined effect of the brood and the queen was larger than each of the effects separately. For example, while the brood did not decrease worker ovary activation and the queen alone only partially decreased it, the combined presence of the queen and the brood fully inhibited ovary activation (Fig. 2b). Similarly, *krh1* levels were more affected by the combined presence of the brood and the queen than by each of them alone (Fig. 4). The synergetic interactions in our study induced physiological changes (ie, ovary activation or *krh1* levels that correlate with oocyte size). This may suggest that costly physiological changes (e.g. workers refraining from activating their ovaries), have a higher threshold for signals to take effect. This could explain why bumble bee worker ovaries are inactive only in young, full-sized colonies (Duchateau and Velthuis, 1988) where both the queen and young brood are present.

Results obtained in experiments examining egg laying, aggressive behavior and *vg* expression levels indicate that some of the effects of the queen and the brood are additive. The brood acted in a manner similar to the queen but to a much smaller extent. For example, while brood caused a two-fold reduction in the *vg* expression, the queen caused a ten-fold reduction in the same transcript (Fig. 4), and while brood significantly reduced worker egg laying, the queen inhibited it completely (Fig. 2a). The additive interactions in our study were typical to behavioral changes that are reversible and thus have a lower threshold for signals to cause a change, resulting in workers responding to either the presence of the queen or the brood, as well as to both. Indeed, not only egg laying and aggression reside under the strict definition of a behavioral change, but also the expression levels of *vg*. While *vg*, a gene typically encoding to the yolk protein invested in female ovaries, is regulated by JH in most insects, it was suggested to decouple from JH in the transition to advanced eusociality and to regulate aggressive behavior in *B. terrestris* (Amsalem et al., 2014b), *B. impatiens* (Padilla et al., 2016) and Themnothorax ants (Kohlmeier et al., 2019; Kohlmeier et al., 2018). In the latter, *vg* was duplicated and its ortholog was associated with behavioral maturation.

In certain cases, the effects of the queen and brood were redundant. This type of interaction is truly puzzling since it questions the need of either the queen or the brood for exerting the full effect. Both queen and brood acted similarly either separately or when combined, as in the case of *mfe* and *dnmt3* expression levels that were equally downregulated in the presence of the queen, the brood, or both (Fig. 4). Levels of *mfe* expression were reduced to the same extent -- twofold in this case – by the queen and the brood and the combination of the queen and the brood did not act any stronger than each of them separately. Furthermore, the brood alone produced a larger effect on the expression levels of *dnmt3* than the queen and the brood together. This suggests that both the queen and the brood probably use the same regulatory lever to affect certain genes, and each of them can exploit the full capacity of that regulatory mechanism. However, both in the case of *mfe* and *dnmt3*, the effects, even though statistically significant, were minor (ca. 1.2-2 fold change) and it is unclear to what extent the queen or the brood utilize these pathways to regulate worker reproduction, and what the underlying mechanism might be.

Our findings on gene expression pattern in response to the queen and the brood contrast previous studies on the honeybee in which both queen and brood pheromones affect worker ovary activation (Mohammedi et al., 1998; Traynor et al., 2014), but no common pattern was observed for the effects of brood pheromone and queen pheromone on worker brain gene expression (Alaux et al., 2009; Grozinger et al., 2003). Previous gene expression studies showed that in the honey bee, *vg* expression is elevated, rather than reduced, following exposure to the queen pheromone, QMP (Fischer and Grozinger, 2008), in line with its role in regulating division of labor in the honeybee, as opposed to regulating reproduction in most other insects. However, the titers of *vitellogenin* protein are reduced in brood tending workers (Amdam et al., 2009; Eyer et al., 2017; Smedal et al., 2009). It further showed that *krh1* levels were reduced following exposure to QMP and were higher in nurses than in foragers (Grozinger and Robinson, 2007), but the precise effect of brood pheromone on honeybee *krh1* expression is still unknown, and *mfe* levels in the honeybee seem to be largely unaffected by exposure to brood (Eyer et al., 2017). Curiously, whole-body levels extracts showed higher *mfe* expression in nurses than in foragers, while in isolated *corpora allata* the opposite was true (Bomtorin et al., 2014; Corona et al., 2019). In our study, however, all of these genes were affected by both the queen and the brood in the same way though to a different extent. This discrepancy suggests that the honey bee, a more derived species, features a larger and more diverse repertoire of regulatory mechanisms than the more primitive species where effects of different social factors use the same focal regulatory levers. The idea that social evolution is characterized with evolutionary diversification of regulatory pathways was proposed for species rather far from one another on the tree of life (e.g. drosophila vs. honeybees) (Robinson and Ben-Shahar, 2002; Toth et al., 2010). Comparison of closely related species exhibiting different eusocial organizations (i.e. bees) would clarify whether the repertoire of regulatory mechanisms has expanded in species exhibiting a stronger reproductive skew.

Our finding that both queen and brood can reduce worker egg-laying, but only queens significantly affect ovary activation, suggests that these two reproductive processes are separately regulated by different physiological and neural pathways. Previous studies in solitary insects demonstrated that oviposition was controlled by distinct neural structures different from those that regulated ovary development, and while the former is more likely under direct innervation control, the latter is subject to neuroendocrine regulation (Meola and Lea, 1972; Mouton, 1971; Thomas and Mesnier, 1973). However, separate mechanisms regulating these processes have not been studied in detail in social insects. Our study highlights the importance of distinguishing between different aspects of reproduction and regulatory mechanisms behind each of them. Ovary activation is a long-term physiological process involving metabolic activity and accompanied by a number of large-scale changes in an organism. Egg-laying, however, is a behavioral phenomenon under CNS control. The fact that brood on its own was capable of affecting egg laying and aggressive behavior but not ovary activation suggests that the effect of brood is limited to behavioral processes but probably does not encompass other pathways regulating ovary activation which the queen can exert influence.

Overall our study sheds light on the synergetic and additive mechanisms of reproductive regulation and maintenance of social harmony in insect societies beyond queen semiochemicals and paves the way to further studies of multiple interacting factors involved in regulating worker reproduction. However, future research is required to understand other factors at play in this system, that remained unexplored in the current study. These include the specific molecular pathways through which the queen and the brood act and the extent to which the queen herself might be influenced by her brood. We hope that our study will blaze the trail for in-depth research of those questions.

## Acknowledgments

We thank members of the Amsalem Lab for helpful discussions and critical reading of earlier drafts of the manuscript and two anonymous reviewers whose comments have greatly improved this manuscript.Captions to figures

