## Supplemental Table S1 for "A small family business: synergistic and additive effects of the queen and the brood on worker reproduction in a primitively eusocial bee"

Table S1. List of genes examined in this study, their accession numbers, and primer sequences

| *Gene* | *Accession Numbers* | *Forward primer* | *Reverse Primer* |
| --- | --- | --- | --- |
| Arginine kinase (HKG) | XM_012391881.2 | GTTGGTAGGGCAGAAGGTCA | AGGTCTACCGTCGTCTGGTG |
| Phospholipase A2 (HKG) | XM_003491149.3 | CATTTCCGCAAGTGGTAGGT | GGTCACACCGAAACCAGATT |
| Vitellogenin (*vg*) | XM_003492229 | CAGCCGCCAATATGATACCT | CCCTCCGTTCGAAGTGATAA |
| Kruppel homologue 1 (*krh1*) | XM_024372425.1; XR_001102792.2; XM_003485596.3 | GAATTGCCAAATCGAGAGGA | GAGGGAGTATGGCATCAGGA |
| Methyl farneosoate epoxidase (*mfe*) | XM_003484680.3 | CAGCCGCCAATATGATACCT | GATCACGCCTGGGTGGTTTA |
| DNA methyltransferase (*dnmt3*3) | XM_012386439.2, XM_024368071.1, XM_024368070.1, XM_024368069.1, XM_024368068.1 | CCTTTTGGATCATAGAGGCCG | ACGTGCAAGATATCACGAAGGA |
